# In-solution Y-chromosome capture-enrichment on ancient DNA libraries

**DOI:** 10.1101/223214

**Authors:** Diana I Cruz-Dávalos, María A Nieves-Colón, Alexandra Sockell, G David Poznik, Hannes Schroeder, Anne C Stone, Carlos D Bustamante, Anna-Sapfo Malaspinas, María C Ávila-Arcos

## Abstract

**Background:** As most ancient biological samples have low levels of endogenous DNA, it is advantageous to enrich for specific genomic regions prior to sequencing. One approach – in-solution capture-enrichment – retrieves sequences of interest and reduces the fraction of microbial DNA. In this work, we implement a capture-enrichment approach targeting informative regions of the Y chromosome in six human archaeological remains excavated in the Caribbean and dated between 200 and 3,000 years BP. We compare the recovery rate of Y-chromosome capture (YCC) alone, whole-genome capture followed by YCC (WGC+Y) versus non-enriched (pre-capture) libraries.

**Results:** We recovered 17–4,152 times more targeted unique Y-chromosome sequences after capture, where 0.01-6.2% (WGC+Y) and 0.01-23.5% (YCC) of the sequence reads were on-target, compared to 0.0002-0.004% pre-capture. In samples with endogenous DNA content greater than 0.1%, we found that WGC followed by YCC (WGC+Y) yields lower enrichment due to the loss of complexity in consecutive capture experiments, whereas in samples with lower endogenous content, WGC+Y yielded greater enrichment than YCC alone. Finally, increasing recovery of informative sites enabled us to assign Y-chromosome haplogroups to some of the archeological remains and gain insights about their paternal lineages and origins.

**Conclusions:** We present to our knowledge the first in-solution capture-enrichment method targeting the human Y-chromosome in aDNA sequencing libraries. YCC and WGC+Y enrichments lead to an increase in the amount of Y-DNA sequences, as compared to libraries not enriched for the Y-chromosome. Our probe design effectively recovers regions of the Y-chromosome bearing phylogenetically informative sites, allowing us to identify paternal lineages with less sequencing than needed for pre-capture libraries. Finally we recommend considering the endogenous content in the experimental design and avoiding consecutive rounds of capture for low-complexity libraries, as clonality increases considerably with each round.

## Background

Uniparental markers such as those on the mitochondrial chromosome (mtDNA) and on Y-chromosome DNA (Y-DNA) are widely used to infer the demographic histories of specific human lineages [1]. Although much smaller than the nuclear genome, the inheritance mechanism and lack of recombination make them powerful tools for inferring ancestry and estimating the ages of pedigrees and the times to the most recent common ancestors (TMRCA) of the mtDNAs and Y-DNAs of present-day populations [2]. Analyses of modern and ancient mtDNA and Y lineages have broadened our knowledge of diversification and founder events from human population history [3, 4, 5].

Due to the large number of copies in each cell, mtDNA has been at the forefront of ancient DNA research [6]. In contrast, each cell possesses just one copy of the Y chromosome. Thus, when analyzing ancient samples, the probability of retrieving any given portion of Y-chromosome DNA is much lower than for mtDNA. Furthermore, as endogenous DNA is often found highly fragmented and in low quantity, recovering ancient DNA (aDNA) from a pool of endogenous and contaminating environmental DNA is extremely challenging and costly. To overcome these challenges, methods have been developed to increase the endogenous DNA proportion of sequencing libraries. These methods target select genomic regions, such as SNPs, whole chromosomes, or mitochondrial or nuclear genomes [3, 7, 8, 9, 10] prior to sequencing. They consequently increase the proportion of genomic regions of interest while reducing sequencing costs.

There are two main types of enrichment methods: solid phase enrichment [11, 12], wherein DNA probes are fixed to a surface, and in-solution enrichment, wherein biotinylated DNA or RNA probes hybridize to the targeted molecules of a DNA library [7, 10, 13]. In the in-solution approach, the probe-target complex is captured with streptavidin-coated magnetic beads, and the remaining fragments, including those from microbial DNA contamination, are washed away. Capture-enrichment approaches have enabled DNA retrieval from samples that initially showed small amounts of endogenous DNA [9, 14, 15]. Consequently, these enrichment methods have positively impacted ancient genomics research by lowering endogenous content requirements, thereby increasing the number of samples that can be employed for profiling.

Capture-enrichment strategies have been applied to target genome-wide SNP sets and to specific subsets of the genome to study the phylogenetic context of ancient populations. Recent implementations [3, 5] include probes targeting thousands of autosomal and Y SNPs characterized by the Simons Genome Diversity Project [16] and the International Society of Genetic Genealogy (ISOGG, https://isogg.org/). However, due to the provenance of the samples of the ISOGG consortium, ISOGG SNPs are best suited to genotype present-day European haplogroups. Consequently, aDNA enrichment has been applied to study Y-chromosome variation in ancient European and Middle-Eastern individuals, while studies of Africans [17] and Native Americans have been restricted either to direct interrogation of known Y-DNA markers with targeted PCR-based sequencing [18] or to low and medium-depth whole-genome sequencing [17, 19, 20, 21].

An important consideration of enrichment designs targeting pre-selected SNPs is the ascertainment bias introduced and the impediment of discovering new variants. An ideal strategy would involve capturing the whole Y chromosome, however its abundance of repetitive sequences makes it less amenable for capture experiments [13]. To overcome these constrains, we made use of a probe design targeting 10.3 megabases (Mb) of the 57.2 Mb Y chromosome defined by Poznik and colleagues [2]. The 41 regions within this 10.3 Mb were selected to fall within the non-recombining portion of the Y chromosome, be depleted of repeats, and well suited for genotype calling from short read sequence data [2]. Furthermore, we were interested in assessing whether Y-DNA could be enriched from libraries with very low endogenous content that had been subjected to WGC and for which we also had pre-capture libraries. We thus tested this approach and compared different enrichment strategies, on samples excavated from the Caribbean, a region that poses a particular challenge for DNA preservation. Previous studies on some of these samples failed to obtain enough Y-DNA data to reliably call a haplogroup even after WGC [22]. Consequently, we investigated the parameters affecting the quality and the quantity of the data and, at the same time, assessed the extent to which the enrichment improved the resolution of the Y-chromosome haplogroup assignment. Our results provide a better understanding of the parameters that affect Y-DNA enrichment experiments and illustrate its benefits for studying the paternal genetic ancestry of ancient human populations.

## Methods

### Samples

We performed 18 capture-enrichment experiments on DNA libraries obtained from the archaeological remains of six individuals excavated from Caribbean contexts. Two samples (STM1 and STM2) belong to 17th-Century enslaved males of African origin from Saint Martin (Lesser Antilles) and were previously reported in [22]. The other four (PI174, PI383, PI435, and PI437) were obtained from archaeological remains from the Paso del Indio site (PI) in Puerto Rico. These four dated between 824 and 1039 CE, as described in [23, 24].

### Ancient DNA extraction

DNA from the STM samples was extracted from tooth roots, as described in [22]. Sampling and DNA extractions for PI samples were conducted at the Arizona State University Ancient DNA Laboratory, a Class 10,000 clean-room facility. Teeth were cleaned with a 1% sodium hypochlorite solution, and the outer surfaces of the tooth roots were mechanically removed with a Dremel tool. Teeth were sliced transversely at the cemento-enamel junction using the Dremel. The roots were then covered in aluminum foil and pulverized by blunt force with a hammer, as in [25]. To avoid contamination, additional precautions were taken, including single use of Dremel wheels, bleach decontamination and UV irradiation of tools and the work area before and between uses, as well as full body coverings for all researchers [26]. DNA was extracted following [14], using 50 mg of pulverized tooth material. Extracts and extraction blanks were quantified with the Qubit 2.0 High Sensitivity assay [27].

### Ancient DNA library preparation

DNA extracted from STM samples was built into 6-bp-indexed double-stranded Illumina libraries, as described in [22]. For PI samples, double-stranded Illumina libraries were prepared following the protocol in [28]. Extraction blanks were also converted into libraries, and an additional negative library control containing only ddH_2_O was also included. 1:100 dilutions of each library were prepared for quality screening through Real-Time PCR (qPCR) using the Thermo Scientific Dynamo SYBR Green qPCR kit with ROX. Reactions were run in triplicate and prepared in final volumes of 20 μl with the following conditions: 10 μl of 2X Dynamo SYBR Green qPCR Master Mix with 0.3 × ROX, 1 μl of primer IS7 (5’-ACACTCTTTCCCTACACGAC-3’) at 10 μM, 1 μl of primer IS8 (5’-GTGACTGGAGTTCAGACGTGT-3’) at 10 μM, 7 μl of ddH_2_O, and 1 μl of library dilution. Reactions were heated to 95°C for 10 minutes for initial denaturation, and further denaturations were performed at 95° C for 15 seconds and for 40 1-minute cycles at 60° C. A final disassociation stage was added at the end of these cycles: 95°C for 15 seconds, 60°C for 15 seconds and 95°C for 15 seconds. Quantification was performed using an ABI7900HT thermocycler and analyzed with SDS software. After qPCR, all libraries were double-indexed as in [29]. To increase library complexity, four 100 μl indexing reactions were performed per library with the following conditions: 10 μl of Pfu Turbo Buffer, 2.50 μl of 10 mM dNTPs, 1.50 μl of 10 mg/ml Bovine Serum Albumin, 2 μl of P5 indexing primer (5’-AATGATACGGCGACCACCGAGATCTACAC xxxxxxACACTCTTTCCCTACACGACGCTCTT-3’) at 10,000 nM, 2 μl of P7 indexing primer (5’-CAAGCAGAAGACGGCATACGAGAT xxxxxxGTGACTGGAGTTCAGACGTGT-3’) at 10,000 nM, 72 μl of ddH_2_O, 1.00 μl of *Pfu* Turbo enzyme (Agilent), and 9 μl of DNA library. Reactions were heated to 95°C for 15 minutes for initial denaturation. Further denaturation, annealing, and elongation were performed at 95°C for 30 seconds, at 58°C for 30 seconds, and for 10 45-second cycles at 72° C. Final extension was performed at 72°C for 10 minutes and reactions were then kept at 10^°^C. All four aliquots of each amplified library were combined, and the library was purified with the Qiagen MinElute PCR purification kit following manufacturer’s instructions with the following modification: the EB buffer was preheated to 65°C before use, and reactions were eluted in 30 μl. A 1-μl aliquot of each library was used for quantification with the Qubit 2.0 Broad Range assay. Purified libraries were further diluted to a factor of 1:1,000 and quantified with the KAPA Library Quantification kit (Kapa Biosystems) following manufacturer’s instructions.

Indexed libraries were amplified a second time to increase the amount of DNA. To increase library complexity, four 100-μl amplification reactions were performed per library. PCR conditions were: 10 μl of 10X Accuprime *Pfx* reaction mix, 3 μl of IS5 primer at 10 μM, 3 μl of IS6 primer at 10 μM, 76 μl of ddH_2_O, 1 μl of Accuprime™ Pfx enzyme, and 7 μl of DNA library. Reactions were heated to 95°C for 2 minutes for initial denaturation, and further denaturation, annealing, and elongation were performed at 95°C for 15 seconds, 60°C for 30 seconds, and for 7–13 1-minute cylces at 68° C. Final extension was performed at 68° C for 5 minutes and reactions were then kept at 4^°^ C. All four aliquots of each amplified library were combined, and the library was purified with Qiagen MinElute PCR purification kit as detailed above. 1 μl of each purified and amplified library were used for flourometric quantification. Purified libraries were further diluted to a factor of 1:10,000 and quantified with the KAPA Library Quantification kit (Kapa Biosystems) following the manufacturer?s instructions. 1 μl of each library was used for fragment analysis with the Agilent 2100 Bioanalyzer DNA 1000 chip.

### Whole-Genome Capture (WGC)

Whole-Genome Capture was performed on each of the libraries obtained from the six archaeological samples (STM1, STM2, PI174, PI383, PI435, PI437) following published protocols. We used the human whole-genome enrichment kit MYbaits (MYcroarray, Ann Arbor, online version 1.3.8) to capture STM libraries, as reported in [15]. For PI libraries, we implemented the WISC approach [9], starting with 500 ng per library and hybridizing for 66 hours. The libraries were PCR amplified for 15–20 cycles.

### Y-chromosome bait design

We used DNA biotinylated probes (baits) from Nimblegen’s SeqCap EZ Choice XL Enrichment Kit for Y capture. Baits were designed using Roche’s NimbleDesign algorithm to target 10.3 Mb well suited for reliable genotype calling and haplogroup inference [2]. The 2.1 million probes were 105 bp in length, with an average tiling overlap of 21 bp for adjacent probes. Probes were designed using the hg19/GRCh37 reference sequence for the Y chromosome, most of which is derived from a single European haplogroup.

### Y-chromosome capture (YCC)

We performed Y-chromosome capture-enrichment experiments on both pre-capture and WGC libraries. Libraries were pooled in equal masses. Capture reactions were performed according to NimbleGen SeqCap EZ XL protocol, with the following modifications: due to limited sample availability, the total mass of the pooled libraries was ^;500 ng rather than the recommended 1.25 μg; hybridization was performed for a total of 65 hours (48–72 h recommended); and the adapter-blocking oligonucleotides were IDT xGen blocking oligos, rather than those sold by Roche, due to increased on-target rates with the IDT xGen oligos. Following capture, libraries were amplified with 6 cycles of PCR, and quality was assessed using the Agilent Bioanalyzer High Sensitivity kit.

### Illumina sequencing

Pre-capture and WGC libraries for the STM samples were sequenced at the National High Throughput DNA Sequencing Centre in Copenhagen, Denmark, on a HiSeq2000 platform using single-end 100-bp runs, as reported in [22] and [15]. PI pre-capture and WGC libraries were paired-end sequenced on the NextSeq500 using a High Output 150-cycle kit with paired-end 76-bp reads. All libraries subjected to YCC (i.e., YCC and WGC+Y libraries from all samples) were sequenced on the NextSeq500 using the High Output 150 cycle kit at Sanford University using paired-end 76-bp mode.

### Sequence data processing and mapping

FASTQ-format reads from the pre-capture and WGC conditions are available for the STM samples through the European Nucleotide Archive, project PRJEB8269, experiment accession numbers ERX682089, ERX682243, ERX682248, and ERX682249 [22]. We processed these reads, as well as the reads generated for this study (YCC and WGC+Y libraries for the STM and all PI libraries) with the following steps. To trim adapters and low quality bases, we used AdapterRemoval v2 with the default options in single-end mode for STM pre-capture and WGC libraries, and in paired-end mode for all PI and STM YCC libraries [30]. As the yield per experiment is variable and can bias the comparisons, we subsampled 10 times an equal number of reads for each experimental condition and for each individual using seqtk (https://github.com/lh3/seqtk). To determine the total number of sequences to subsample, we selected the lowest number of reads that passed the trimming filters for each individual across experiments (Table 1). The sequences were then aligned to the Homo sapiens reference genome build 37 (hg19) using the BWA aligner [31] implemented in PALEOMIX [32], with a mapping quality threshold set at 30. The quality of the aligned bases was rescaled with mapDamage2 [33] to lower the quality of mismatches to the reference sequence that likely derive from DNA damage.

**Table 1.** Total number of reads. Ten replicates per library were obtained by down-sampling to the minimum number of retained reads within each sample (underlined). The “Mapping to chrY”and “% of on-target”sections indicate the number of unique reads mapping to the Y-chromosome, and the percentage of unique on-target reads respect to the total reads mapping to the Y-chromosome, respectively.

|  |  | STM1 | STM2 | PI174 | PI383 | PI435 | PI437 |
| --- | --- | --- | --- | --- | --- | --- | --- |
| Pre-capture | Total reads | 34,025,874 | 14,973,474 | 9,173,100 | 2,795,632 | 4,617,394 | 9,986,135 |
|  | mapping to chrY | 1,384 | 301 | 4 | 3 | 6 | 6 |
|  | % of chrY on-target | 74% | 72% | 25% | 67% | 50% | 33% |
| YCC | Total reads | 68,795 | 41,884 | 159,728 | 107,796 | 41,810 | 56,157 |
|  | mapping to chrY | 16,430 | 2,562 | 17 | 27 | 26 | 12 |
|  | % of chrY on-target | 99% | 99% | 100% | 100% | 96% | 100% |
| WGC | Total reads | 14,973,474 | 29,884,294 | 10,052,798 | 10,265,430 | 12,363,127 | 9,132,306 |
|  | mapping to chrY | 6,407 | 1,925 | 5 | 8 | 57 | 7 |
|  | % of chrY on-target | 62% | 63% | 60% | 100% | 65% | 57% |
| WGC+Y | Total reads | 98,540 | 30,629 | 17,212,495 | 14,557,575 | 10,181,433 | 11,356,277 |
|  | mapping to chrY | 4,854 | 236 | 37 | 152 | 242 | 47 |
|  | % of chrY on-target | 96% | 95% | 100% | 99% | 97% | 96% |

To calculate the enrichment rate, we used the subsampled data and computed the average number of unique reads mapped to the on-target regions in each experiment. Then, we calculated fold-enrichment by dividing the on-target average of YCC or WGC+Y experiments by that of the pre-capture libraries. Specifically, we calculated this fold-enrichment for YCC using the pre-capture condition as a baseline and compared the WGC+Y experiments to both the pre-capture and WGC conditions. When we observed no reads aligning to the Y chromosome or target regions, we used the maximum number of reads observed across the replicates of a given library as a baseline. We computed binomial proportion confidence intervals for the mean endogenous content, the proportion of on- and off-target reads, and clonality, and we conducted a t test for the length estimate. Confidence intervals and chi-squared tests were computed in R software, version 3.3.1 [34].

To call Y-chromosome genotypes for each sample, we first merged data across experiments. We used the haploid genotype caller implemented in ANGSD, retaining only bases with quality scores of at least 13 and sampling one random base at each site [35]. Finally, we performed a binary tree search with a custom script to find the most derived SNP that determines the haplogroup of the individuals. We used as input the phylogenetic tree constructed from the Y-SNPs reported in Phase 3 of the 1000 Genomes Project [4].

### Sex determination

To determine the biological sex of the six individuals, we used the script in [36] to calculate the ratio (R_y_) of reads mapping to the Y-chromosome to those mapping to both sex chromosomes [36]. R_y_ values above 0.075 are consistent with a male genotype.

### Yield and enrichment curves

We estimated the yields and complexities of the libraries, with respect to the reads mapping to the targeted regions, with the preSeq package implemented in R (preseqR, [37]) and corrected the amount of required sequencing by the fraction of on-target reads in the libraries. Since the method relies on having a fraction of duplicated reads to estimate the yield, for the cases where the pre-capture libraries did not have duplicated on-target reads to adjust a yield curve, we instead assumed a linear relationship with a slope equal to the proportion of unique on-target reads present in the library. We then modeled an “enrichment curve”to explore the level of enrichment predicted for different amounts of sequencing. To this end, we used the median unique on-target reads estimated by PreSeq to calculate an expected fold-enrichment. We divided the median value estimated by PreSeq of each captured library by the median of its pre-captured counterpart (i.e., YCC vs. pre-capture, WGC+Y vs. pre-capture, and WGC+Y vs. WGC).

## Results

### Enrichment rates

We tested the performance of Y-chromosome capture on Illumina sequencing libraries obtained from the archaeological remains of six individuals excavated in the Caribbean islands of Saint Martin (STM1 and STM2) and Puerto Rico (PI174, PI383, PI435, and PI437) (Table 1). For each sample, we performed a series of enrichment experiments, as depicted in Figure 1. First, we shotgun-sequenced a DNA library without performing any enrichment. We then performed a capture reaction targeting a set of DNA probes covering 10.3 Mb of the non-recombining portion of the Y chromosome. These regions were validated by Poznik and colleagues in [2] as being well suited for unambiguous read mapping and for yielding reliable genotype and haplogroup calls from short-read sequencing. Additionally, we performed another set of capture experiments, either enriching only the whole-genome (WGC) or the on-target regions after having enriched the whole genome (WGC+Y). First, we confirmed the molecular sex of the samples and determined that the six individuals each had a karyotype consistent with XY [36]. We then assessed the performance of the capture experiments.

**Figure 1.**
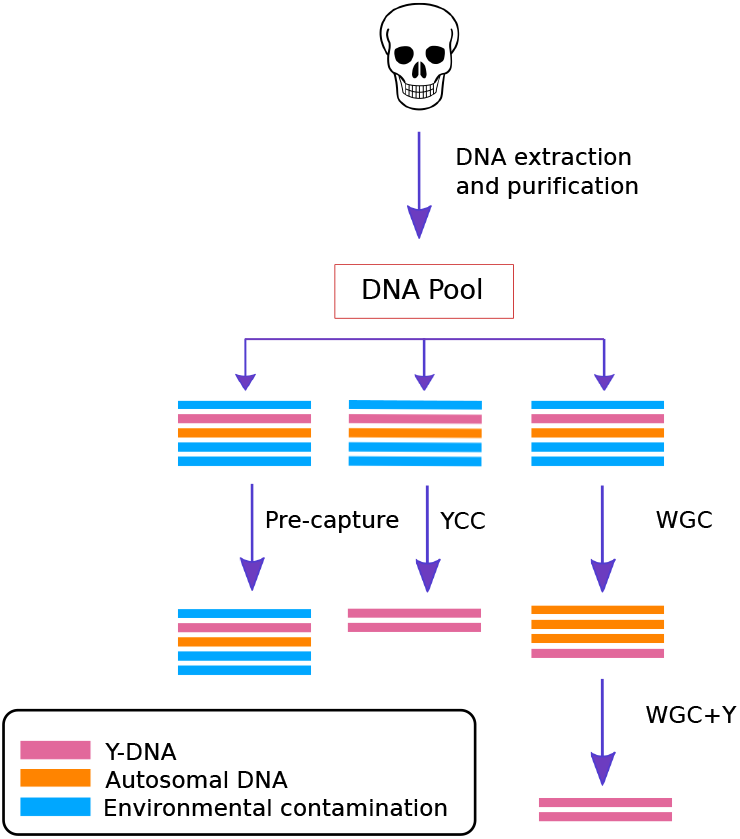
Experimental enrichment scheme. We have four different conditions: Pre-capture, YCC, WGC, and WGC+Y. The pre-capture condition is our initial library preparation prior to any enrichment. The WGC is designed to target all autosomal and sex chromosomes. The Y-capture in the YCC and the WGC+Y conditions targets ~10.3 Mb of Y-chromosome sequence.

Since experiments yielded differing numbers of reads per sample, we down-sampled to equal numbers per individual as described in Methods (Table 1). The pre-capture libraries yielded just 0.01% to 1.54% unique reads aligning to the human genome (Figure 2), and less than 0.004% mapping to the Y-chromosome. After implementing the YCC and WGC+Y enrichments, the endogenous DNA content increased, on average, by factors of 24.2 to 122.0 for the STM samples (Figure 2A) and by factors between 3.2 and 38.9 for the PI samples (Figure 2B). Moreover, in the YCC and WGC+Y libraries, 5.3% to 68.3% of the human STM reads mapped to the Y chromosome, with 17.7% to 50.0% the corresponding figures for the PI samples.

**Figure 2.**
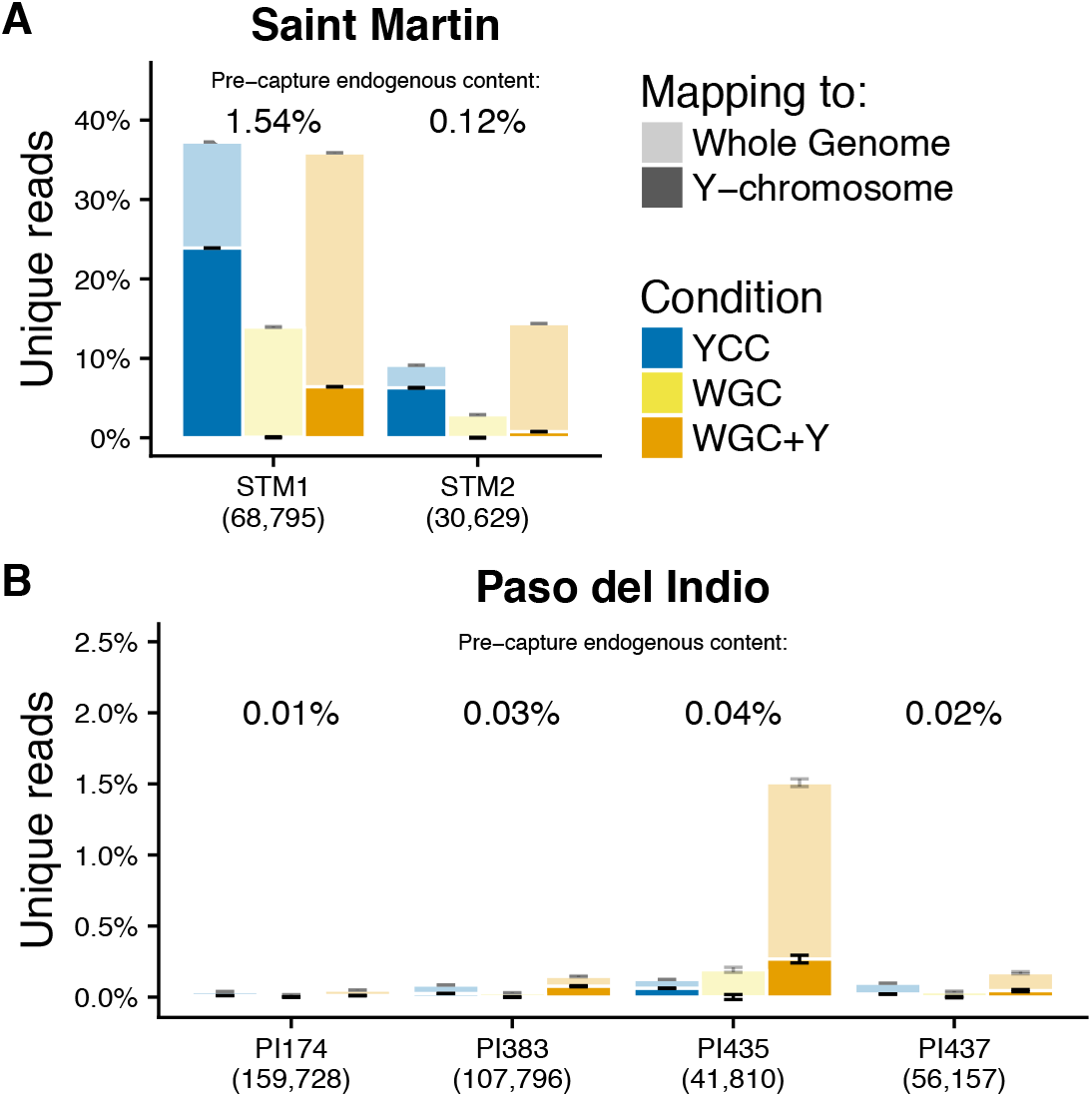
Endogenous DNA content in enriched libraries. Percentage of the unique retained reads that aligned to the human genome in (A) Saint Martin and (B) Puerto Rico samples. “STM”stands for Saint Martin and “PI”for Paso del Indio, Puerto Rico. The percentages in parentheses below the x-axis indicate the number of down-sampled reads per library. The error bars represent 95% confidence intervals of endogenous DNA content found in the samples across the 10 down-sampled replicates. Darker colors correspond to the proportion of the unique reads that aligned to the Y chromosome. Whole-genome enriched libraries have j 0.04% reads aligning to the Y chromosome.

To evaluate whether the enrichment experiments effectively recovered the targeted regions, we compared the total number of unique reads mapping to the targeted regions to the number of off-target reads (Figure 3). YCC experiments on the STM samples yielded 6,476 to 9,549-fold increases of on-target sequences compared to the pre-capture condition. The WGC+Y experiments on the same samples resulted in 105- to 176-fold-enrichment compared to WGC alone. For the PI samples, YCC experiments resulted in 12- to 250-fold enrichment, and we observed 85.5- to 813-fold enrichment for the WGC+Y approach. Although we observed an increase in the off-target content for the STM enrichments (Figure 3), it is one order of magnitude smaller than their respective on-target enrichment. Above 96% of the reads mapping to the Y chromosome were on target in both YCC and WGC+Y, in contrast to the pre-capture and WGC experiments where these figures range between 25% and 72% (Table 1). Overall, the distribution of the on-target sequences in all Y-chromosome enrichment experiments is qualitatively even (Figure 4, Supplementary figure 1). In summary, all Y-chromosome capture enrichment experiments consistently increased the number of unique on-target reads.

**Figure 3.**
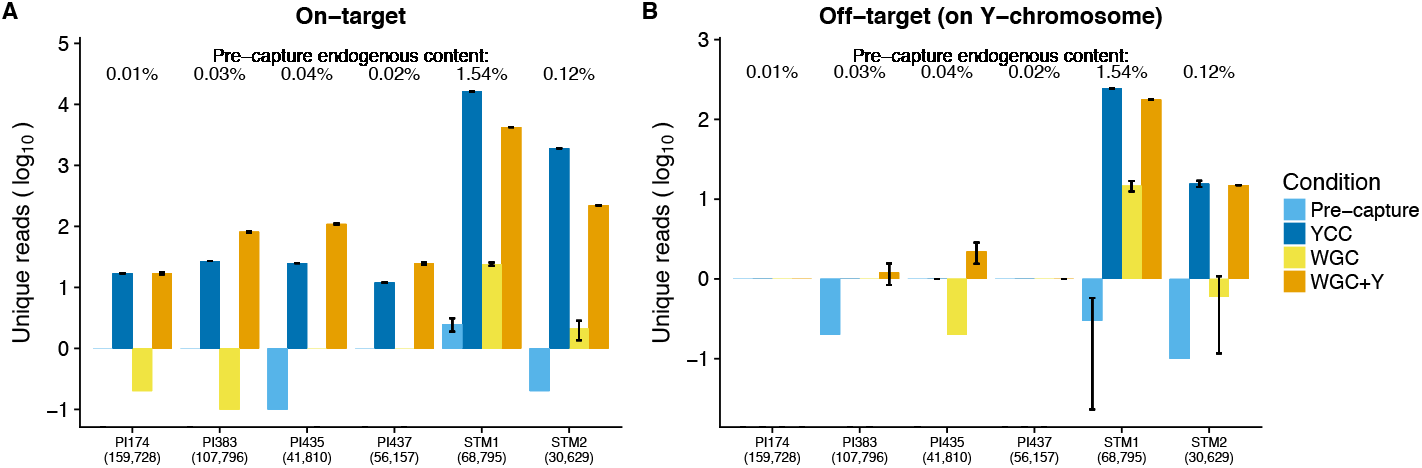
On- and off-target reads. The bars represent the 95% confidence interval for 10 replicates. (A) Unique reads mapping to the Y-chromosome target regions. (B) Unique reads mapping to the Y chromosome but not to the targeted regions. For some libraries, no reads mapped to the Y chromosome, across the 10 replicates.

**Figure 4.**
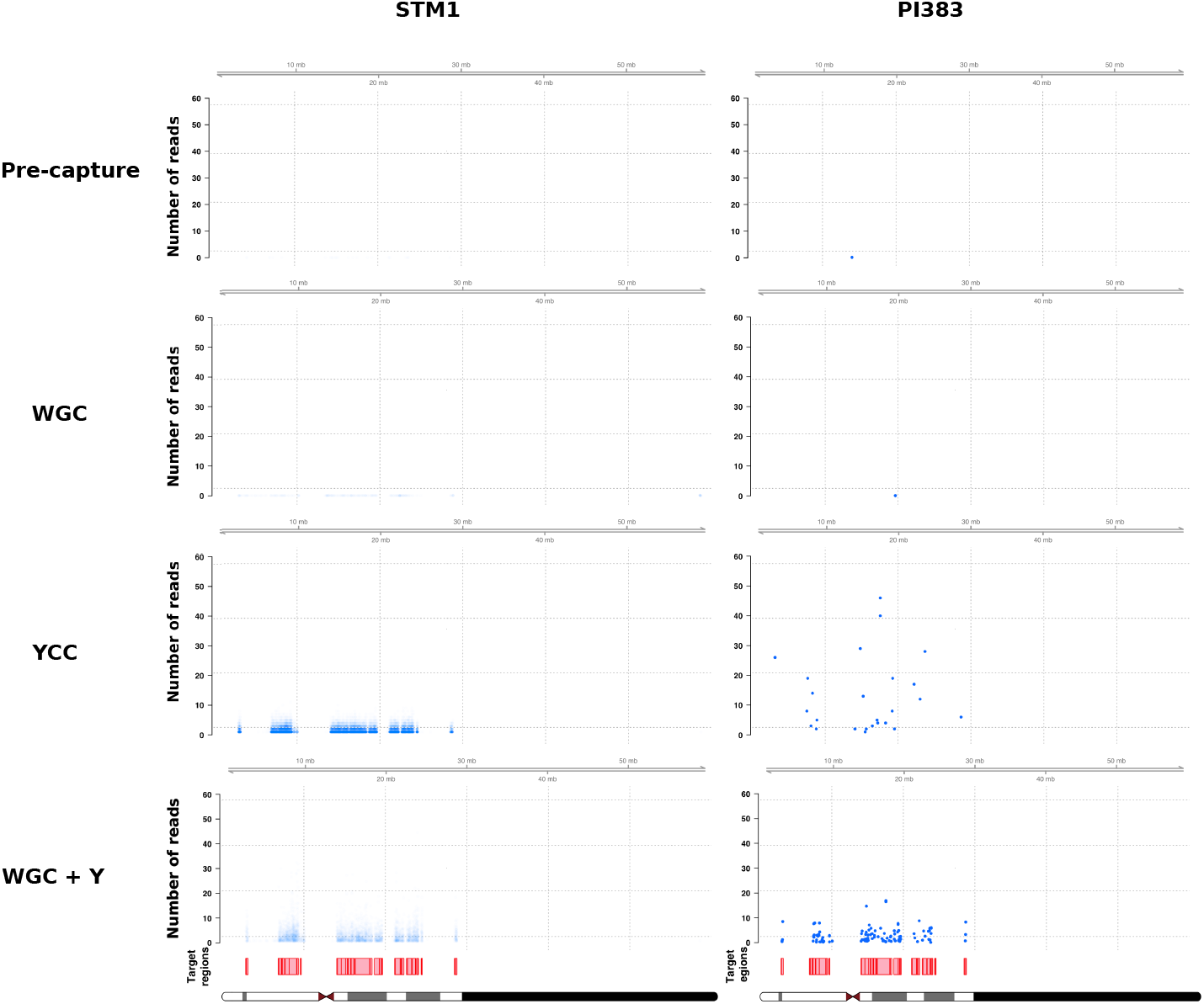
Depth of coverage across the Y-chromosome. From top to bottom, rows depict the coverage levels for the pre-capture, YCC, WGC and WGC+Y conditions. Red boxes represent the targeted regions. Each blue point represents sequencing coverage within a 1,000-bp window, averaged across 10 subsampled replicates per sample per condition explaining depths of coverage below 1. For visual purposes, we increased the opacity of the points in the PI383 column.

### Length distribution and clonality

To explore which features of the pre-capture libraries may have influenced the differences in enrichment rates between the STM and PI groups, we contrasted the lengths and complexities of the individual libraries across experiments. We observed a significant trend toward longer fragments after enrichment for all experiments (Figure 5, Supplementary figure 2) (paired samples t-test, p-value = 0.002), consistent with previous findings [7, 15, 38]. Reads from the STM samples were 91.7 to 92.1 base pairs (bp) long for the pre-capture condition, 87.4 to 94.5 bp after YCC, and 105.5 to 108.3 long after WGC+Y. Likewise, whereas the average length of PI reads in the pre-capture libraries ranged from 63.6 to 69.1 bp, average lengths increased to 76.3–102.3 and 69.3–82.2 and bp after YCC (paired samples t-test, p-value = 0.02) and WGC+Y (paired samples t-test, p-value = 0.03), respectively. On-target clonality levels (percentage of PCR duplicates) were considerable for all samples in the YCC and WGC+Y experiments. We observed 7.0% to 28.6% clonality for YCC of the STM samples at 68,795 and 30,629 down-sampled reads. The remaining WGC+Y (STM), and all WGC+Y and YCC (PI) libraries had greater clonality values, ranging from 65.4% to 94.1% (Figure 6). Although we did not observe on-target duplicates with which to calculate the clonality in any of the down-sampled pre-capture libraries (Supplementary table 2), using the whole data we estimated clonality values ranging from 0.8% to 3.5% (for 101,373,044 and 10,506,505 sequenced reads, Supplementary table 1).

**Figure 5.**
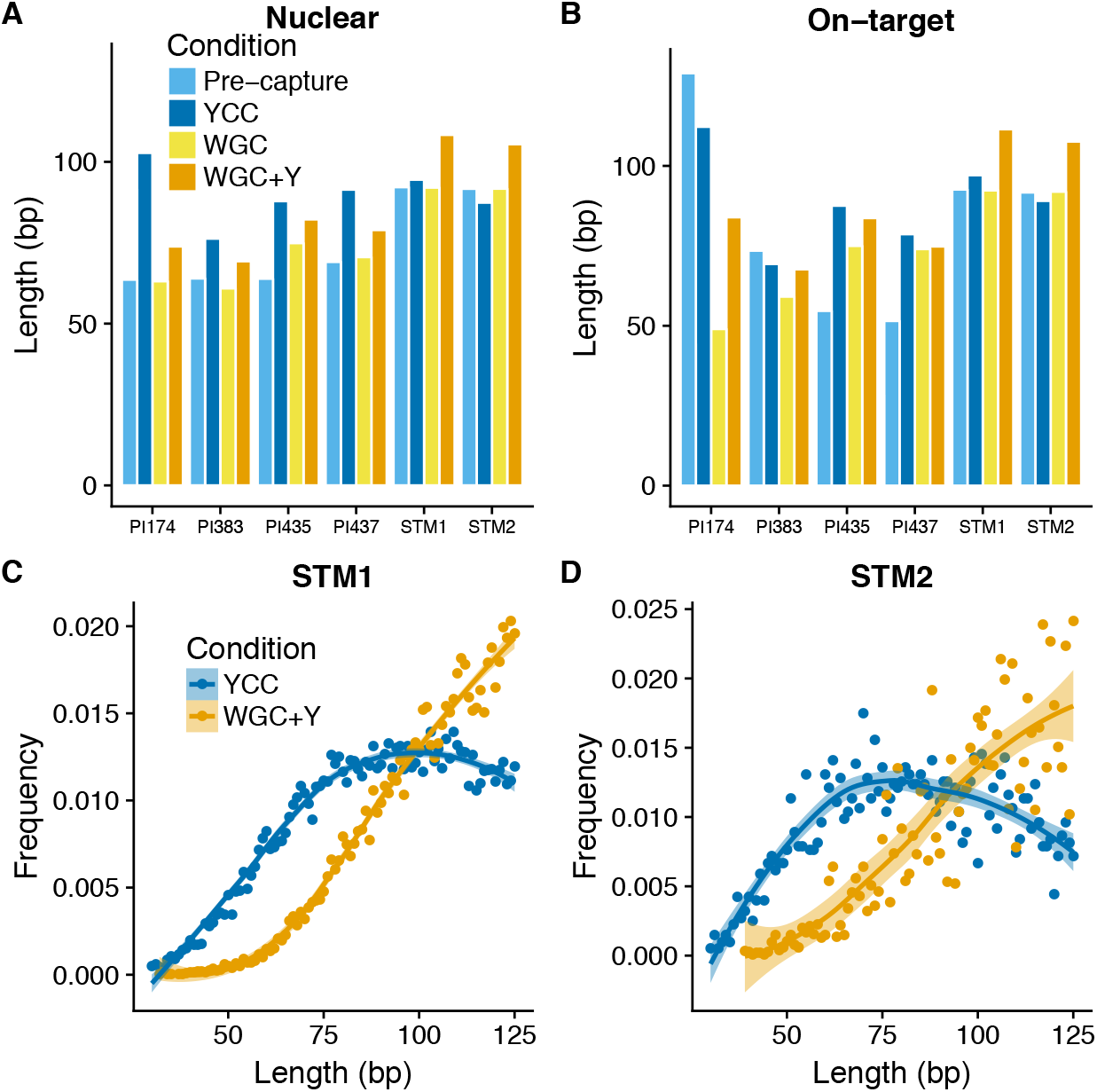
Lengths of mapped reads. (A) Reads aligned to the nuclear genome. (B) On-target reads. (C) and (D) depict the length distributions of reads mapping to the whole genome for STM1 and STM2 samples, respectively. The length distribution was smoothed by fitting a polynomial curve to the observed frequencies; the ribbons correspond to 95% confidence intervals.

**Figure 6.**
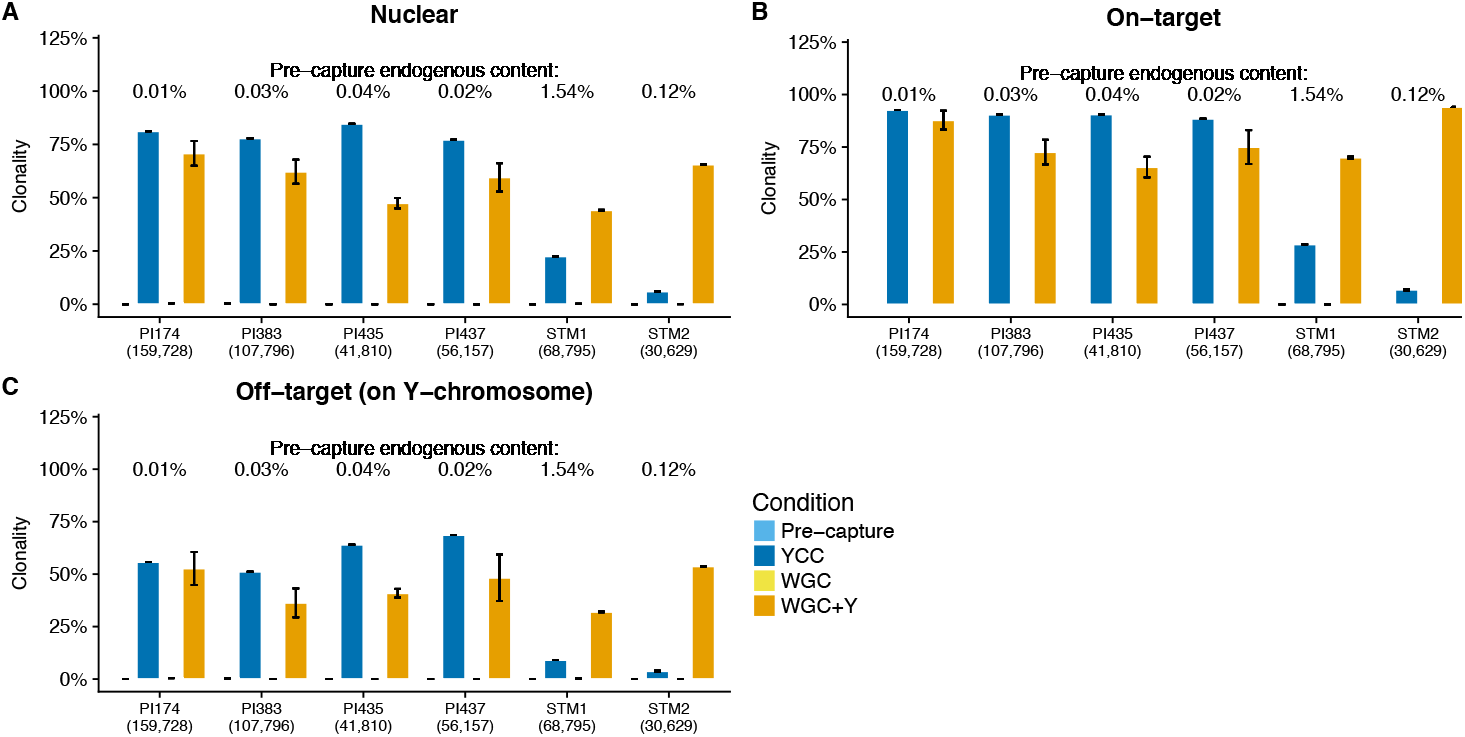
Clonality. (A) Clonal reads mapping to the nuclear human genome. (B) Clonality of the reads mapping to the targeted regions. (C) Clonal reads mapping to the Y chromosome but not to the targeted regions. Error bars represent 95% confidence intervals across 10 subsampled replicates.

### On-target yield for aDNA libraries

The yield curves for both capture conditions (YCC and WGC+Y) corroborate the high clonality of the PI libraries (Figure 7A, B). We observed that the YCC libraries of these samples plateaued at very shallow sequencing, saturating at 25,000 sequenced reads, compared to saturations at 50,000–100,000 sequenced reads for the WGC+Y libraries. However, the complexity curves indicate that after sequencing 100,000 reads of the WGC+Y libraries, we would not retrieve more than 150 different reads, regardless of the capture approach. On the other hand, although we sequenced fewer than 100,000 reads for each of the YCC experiments on the STM individuals, the complexity curves suggest that these libraries could be further sequenced to increase the coverage of the targeted regions.

**Figure 7.**
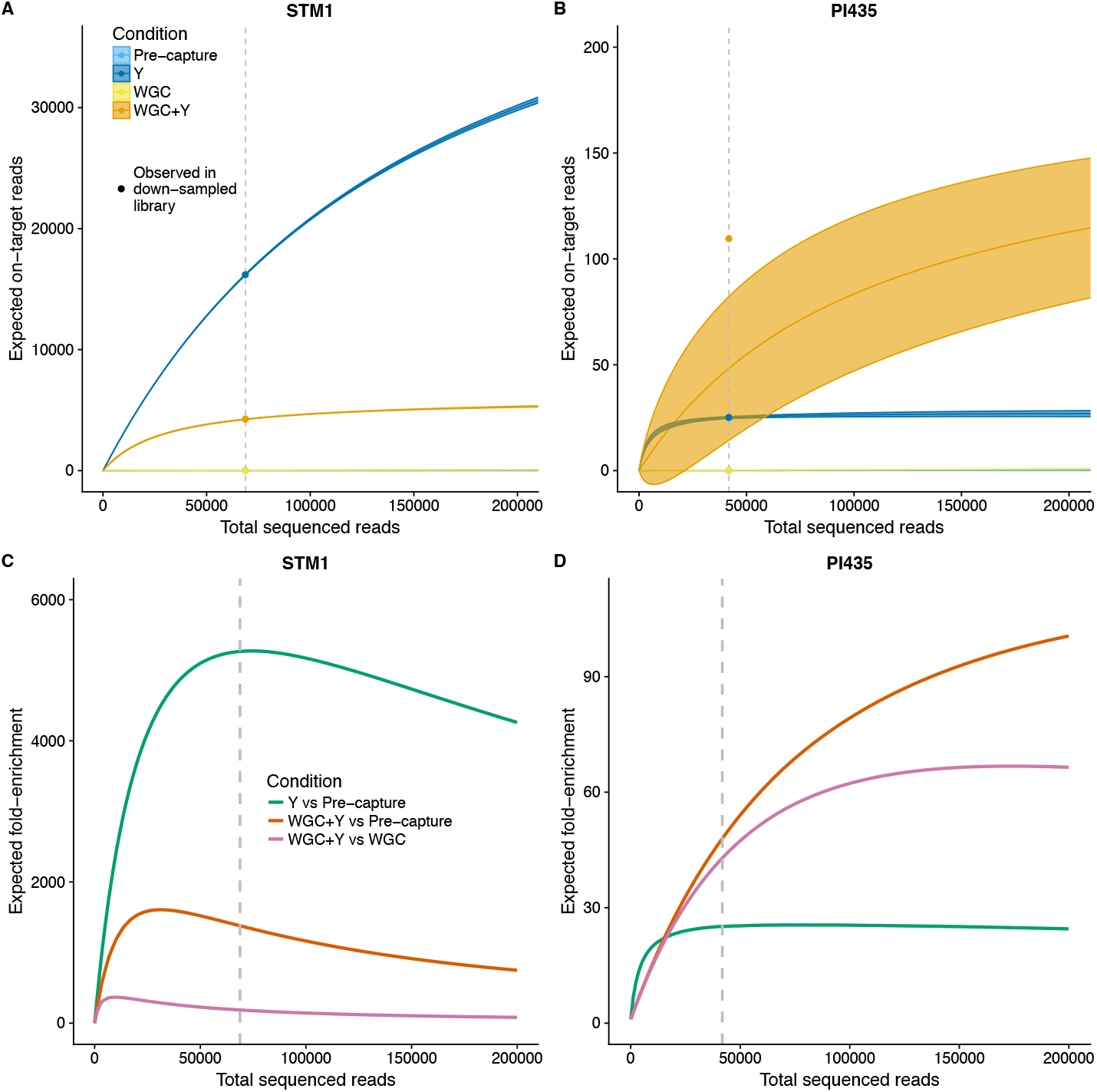
Expected yield and on-target fold-enrichment. Dashed lines indicate the number of down-sampled reads. (A) and (B): Predicted median value and variance (across 100 bootstrap replicates) of the number of on-target reads, as a function of total sequenced reads. The points depict the observed numbers of on-target reads in the down-sampled libraries. (C) and (D): Expected enrichment of on-target reads versus number of sequenced reads for each condition and each sample.

### Enrichment rates based on yield estimates

We computed enrichment curves (Figure 7 C, D) to predict enrichment rates for deeper sequencing. Consistently, we observe that the YCC of STM2 recovers 1,790 more on-target reads than the pre-capture experiment at the down-sample point, but, as it has not reached saturation, this enrichment value can increase to at least 5,000-fold, similar to the maximum enrichment of STM1. In contrast, the projected enrichment of the PI WGC+Y libraries at 200,000 total reads is only 100- and 72-fold versus the pre-capture and WGC libraries, respectively. We also note that, for PI samples, the three enrichment rates calculated “decelerate”at 100,000 sequenced reads or fewer, once more reflecting the low complexity levels of the initial pool.

### Y haplogroup calls

Finally, we combined all the sequence data generated for each sample to call Y-chromosome haplogroups. With the new data generated, we were able to call haplogroups for both STM individuals. The combined on-target depth of coverage was 0.03x to 0.24x, with a sequencing depth of at least one for 0.36–1.96 Mb (Table 2). In the STM1 individual we observed derived alleles belonging to the R1b-M343 clade, consistent with its previously reported haplogroup (R1b1c-V88) [22]. In the same study, even after WGC, the haplogroup for STM2 could not be resolved [22]. Only after integrating the previously generated data with that produced in this study, were we able to assign the Y-haplogroup as E1b1a1a1-M80. Regarding the PI samples, we did not find any reads covering the SNPs in the database in the pre-capture or the WGC libraries. After YCC or WGC+Y, we observe between 5 and 74 variant sites per library, leading to the identification of haplogroups defining broader regions. For instance, the haplogroup found for PI383, P-M45, is the immediate ancestor of haplogroups R and Q. In Table 2, we show the haplogroups for all the individuals according to the most derived SNP identified in each condition.

**Table 2.**
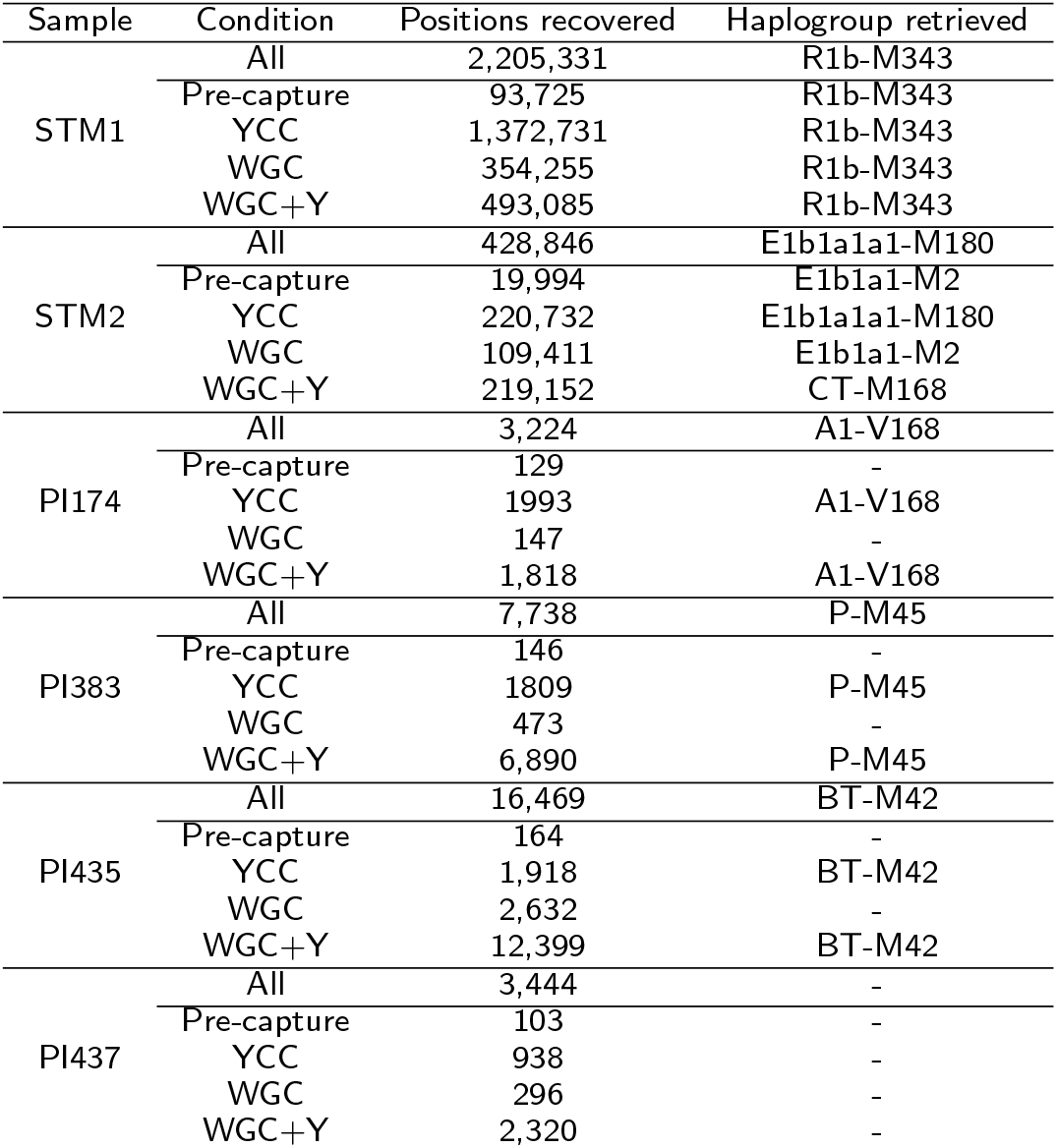
Numbers of Y-chromosome bases retrieved and haplogroups.

## Discussion

In this paper, we have described the efficiency of Y-DNA recovery from in-solution Y-chromosome capture-enrichment experiments and from WGC followed by Y-chromosome enrichment on aDNA libraries obtained from the archaeological remains of six males excavated in the Caribbean and dating between 300 and 2,000 years old. The experimental design involved the targeted enrichment of 10.3 Mb of the Y chromosome on both standard and WGC libraries. As several aspects of the library preparation and WGC enrichment protocols differed between sample groups, we evaluated the success of the enrichment using within-sample comparisons. Overall, both approaches succeed in increasing the proportion of on-target sequences as compared to pre-capture libraries. For both enrichment approaches and for all samples, we observed that most of the sequenced reads mapped to the targeted regions.

### Factors influencing enrichment

Although we observed consistent enrichment of Y-DNA on aDNA libraries, the marked differences of the performance between samples (STM and PI) and library types (standard vs. WGC libraries) is mostly like due to differences in starting endogenous content, read-length distributions, and the complexities of the libraries. Although it is now possible to increase the endogenous content of poorly preserved tissues, it has been shown that in-solution capture enrichment techniques perform better on samples with starting endogenous content greater than 1% and with little clonality [15]. Thus, as expected, the PI sample enrichment levels were systematically lower than those in the STM samples. Low complexity levels in the starting libraries also hamper the success of capture experiments, as these protocols usually involve an amplification step, which further increases the clonality. In addition, the enriched libraries were subject to an increase in the fragment length most likely driven by the probe length (105 bp) (Figure 5). However, the shift was more pronounced for the PI samples, indicating that a substantial proportion of the shorter fragments in the pre-capture and WGC libraries was not retrieved in the YCC and WGC+Y. Together, these observations provide useful insights as to the features in standard libraries that should be considered when planning capture experiments, thereby opening avenues for investigating ways to optimize these protocols.

Subsequently, we noted that for the STM samples the YCC experiments yielded higher fold-enrichments of the targeted regions than did the WGC+Y captures. In contrast, for the PI samples, the WGC+Y performed betted. While we believed the differences owe mostly to the endogenous content, it could also be in part due to the techniques employed to enrich the whole genomes of the STM and PI samples (MYbaits and WISC, respectively). Indeed, although based on the same molecular principle, those techniques have slight differences in their performances [15]. However, despite the slightly higher enrichment rates in WGC+Y libraries, as compared to YCC, for the PI libraries, we do not recommend implementing consecutive enrichments, as we show here that doing so considerably reduces the complexity of the libraries [39] and, consequently, the ability to identify informative SNPs for haplogroup assignment.

### Implications of Y-haplogroups assignments for the samples

Finally, the haplogroup inference was effective only for the two STM samples, for which we recovered at least 0.36 Mb of the Y-chromosome (3.5% of on-target regions). The STM1 individual bore the M343 mutation characteristic of haplogroup R1b, consistent with the back-to-Africa R1b1c-V88 haplogroup reported in [22]; however, we did not observe any of the SNPs specific to the V88 branch. Whereas for STM2, we identified a common and widespread African haplogroup characteristic of the Bantu expansion [40], E1b1a1a1-M80, consistent with the results from the analyses of the autosomal chromosomes [22]. For the remaining individuals, we could not resolve haplogroups due to the low depth at which they were sequenced due to limited budget, although our results might be impacted by the paucity of Y-SNPs that define the tips of the Native American haplogroups versus haplogroups from other well-characterized populations in the database employed. For example, no Puerto Rican males in the 1000 Genomes dataset bore Native American Y haplogroups. Rather, all possessed European or African lineages, primarily belonging to R1b and E1b clades [4], reflecting sex-biased admixture patterns during European colonization of the island (and reproduced across the Americas) [41]. This fact highlights the need to assay ancient genetic variation among pre-contact Native American samples, for which methods such as ours can be of important advantage. It is often challenging to recover DNA for such samples and the enrichment method we discuss here would certainly help in those cases.

## Conclusions

In the past decade, new technologies and protocol improvements have emerged to efficiently recover ancient DNA. However, the endogenous DNA fraction continues to be a limiting step in ancient genomics studies. The first efforts to overcome this limitation have focused on targeting the mtDNA, because it is relatively short (^16 kB), and it is present in multiple copies per cell, unlike the autosomes (two copies) and the Y chromosome (one copy). For the Y chromosome, targeted enrichment strategies are more problematic due to its richness in repetitive and palindromic sequences. For the same reasons, Y-chromosome content is relatively poor in WGC studies, although WGC is becoming a cost-effective alternative for ancient genomics. Therefore, we used previously reported high-quality regions to capture the most phylogenetically informative portion of the Y chromosome. We confirmed the effectiveness of the method by noting that, after capture, up to 99.1% of the reads mapping to the Y chromosome fall within the targeted regions. We observed that YCC and WGC+Y libraries outperformed pre-capture libraries with respect to Y-DNA content. Whether it is advantageous to first perform WGC before enriching for Y-DNA seems to depend on the level of endogenous DNA content, the complexities and fragment lengths of the starting libraries. In this study, libraries with endogenous DNA content greater than 0.12% yielded ^10-fold greater enrichment rates under YCC, as compared to WGC+Y libraries, whereas in the four samples with low endogenous DNA proportions (0.01% to 0.02%), we observed a greater enrichment for WGC+Y experiments, but at the cost of higher clonality. We thus stress the need to consider the initial complexity, endogenous DNA content, and read lengths when planning these experiments. We recommend a design that includes the estimation of predictive yield and enrichment curves, based on shallow sequencing, to inform the best sequencing strategy and avoid sequencing beyond saturation.

Finally, there is a vast potential to incorporate Y-chromosome information from aDNA samples into the study of human population history from regions beyond Eurasia. In our work, we go beyond SNP capture and present the first instance of Y-chromosome capture on ancient samples, opening new avenues of research to improve the performance of these experiments and to extract Y-chromosome information from ancient samples.

## Availability of data and material

YCC and WGC+Y reads of STM samples are available in the European Nucleotide Archive under the accession number PRJEB23498. Y-chromosome alignments for Paso del Indio samples are available in the NCBI Short Read Archive (SRA) under BioProject PRJNA419010.

## Author’s contributions

MCAA conceived the project with input from CDB, DP and AS. MNC, AS and HS performed laboratory work. ASM, DICD and MCAA designed the data analysis strategy. DICD performed most data analyses with input from DGP. DICD and MCAA wrote the manuscript with input from all the authors. All authors read and approved the final manuscript.

## Acknowledgements

Calculations were performed on UBELIX (http://www.id.unibe.ch/hpc), the HPC cluster at the University of Bern. MNC would like to thank William J. Pestle, L. Antonio Curet and Edwin Crespo-Torres for providing the Paso del Indio samples and Meredith Carpenter, Morten Rasmussen and Rosa Fregel for assistance with computational analyses.

## Additional Files

Supplementary table 1. Summary of sequenced and mapped reads of the complete dataset Supplementary table 2. Mean and standard errors regarding reads from down-sampled libraries.

Supplementary figure 1. Depth of coverage across the Y-chromosome.

From top to bottom, rows depict the coverage levels for the pre-capture, YCC, WGC and WGC+Y conditions. Red boxes represent the targeted regions. Each blue point represents sequencing coverage within a 1,000-bp window, averaged across 10 subsampled replicates per sample per condition, explaining depths of coverage below 1. For visual purposes, we increased the opacity of the points in the PI samples.

Supplementary figure 2. Length distribution of mapped reads.

Length distributions of reads mapping to the whole genome. The length distribution was smoothed by fitting a polynomial curve to the observed frequencies; the ribbons correspond to 95% confidence intervals.

Supplementary figure 3. Expected yield and on-target fold-enrichment.

Dashed lines indicate the number of down-sampled reads. (A-F): Predicted median value and variance (across 100 bootstrap replicates) of the number of on-target reads, as a function of total sequenced reads. The points depict the observed numbers of on-target reads in the down-sampled libraries. (G-L): Expected enrichment of on-target reads versus number of sequenced reads for each condition and each sample.

